# A Cdk9-PP1 kinase-phosphatase switch regulates the elongation-termination transition of RNA polymerase II

**DOI:** 10.1101/190488

**Authors:** Pabitra K. Parua, Gregory T. Booth, Miriam Sansó, Bradley Benjamin, Jason C. Tanny, John T. Lis, Robert P. Fisher

## Abstract

The end of the RNA polymerase II (Pol II) transcription cycle is strictly regulated to ensure proper mRNA maturation and prevent interference between neighboring genes^1^. Pol II slowing downstream of the cleavage and polyadenylation signal (CPS) leads to recruitment of cleavage and polyadenylation factors and termination^2^, but how this chain of events is initiated remains unclear. In a chemical-genetic screen, we identified protein phosphatase 1 (PP1) isoforms as substrates of human positive transcription elongation factor b (P-TEFb), the cyclin-dependent kinase 9 (Cdk9)-cyclin T1 complex^3^. Here we show that Cdk9 and PP1 govern phosphorylation of the conserved transcription factor Spt5 in the fission yeast *Schizosaccharomyces pombe*. Cdk9 phosphorylates both Spt5 and a negative regulatory site on the PP1 isoform Dis2^4^. Sites phosphorylated by Cdk9 in the Spt5 carboxy-terminal domain (CTD) are dephosphorylated by Dis2 *in vitro*, and Cdk9 inhibition *in vivo* leads to rapid Spt5 dephosphorylation that is retarded by concurrent Dis2 inactivation. Chromatin immunoprecipitation and sequencing (ChIP-seq) analysis indicates that Spt5 is dephosphorylated as transcription complexes traverse the CPS, prior to or concomitant with slowing of Pol II^5^. A Dis2-inactivating mutation stabilizes Spt5 phosphorylation (pSpt5) on chromatin, promotes transcription beyond the normal termination zone detected by precision run-on transcription and sequencing (PRO-seq)^6^, and is suppressed by ablation of Cdk9 target sites in Spt5. These results support a model whereby the transition of Pol II from elongation to termination is regulated by opposing activities of Cdk9 and Dis2 towards their common substrate Spt5—a bistable switch analogous to a Cdk1-PP1 module that controls mitotic progression^4^.

---

In metazoans and fission yeast, commitment to and exit from mitosis depend on inhibitory phosphorylation of PP1 by Cdk1 and its reversal, respectively^4^. In human extracts, analogue-sensitive (AS) Cdk9 modifies two isoforms of PP1, PP1β and PP1γ, on conserved, carboxy-terminal sites analogous to the PP1α residue labeled by Cdk1^3,7^. Of the two PP1 isoforms in fission yeast, Dis2 and Sds21, only Dis2 has the potential for inhibition by CDKs through phosphorylation of its Thr316 residue (Fig. 1a)^8,9^. Purified S. *pombe* Cdk9 phosphorylated Dis2 but not Sds21 *in vitro* (Fig. 1b); labeling was diminished by a *T316A* mutation changing Thr316 to alanine, but not by a Dis2-inactivating mutation^10^. Moreover, treatment of asynchronously growing *cdk9*^*as*^ but not *cdk9*_+_ cells with the bulky adenine analogue 3-MB-PP1, a selective inhibitor of AS Cdk9^11^, decreased Thr316 phosphorylation of chromatin-associated Dis2 (Fig. 1c), indicating that Dis2 is indeed regulated by Cdk9 *in vivo*.

**Figure 1.**
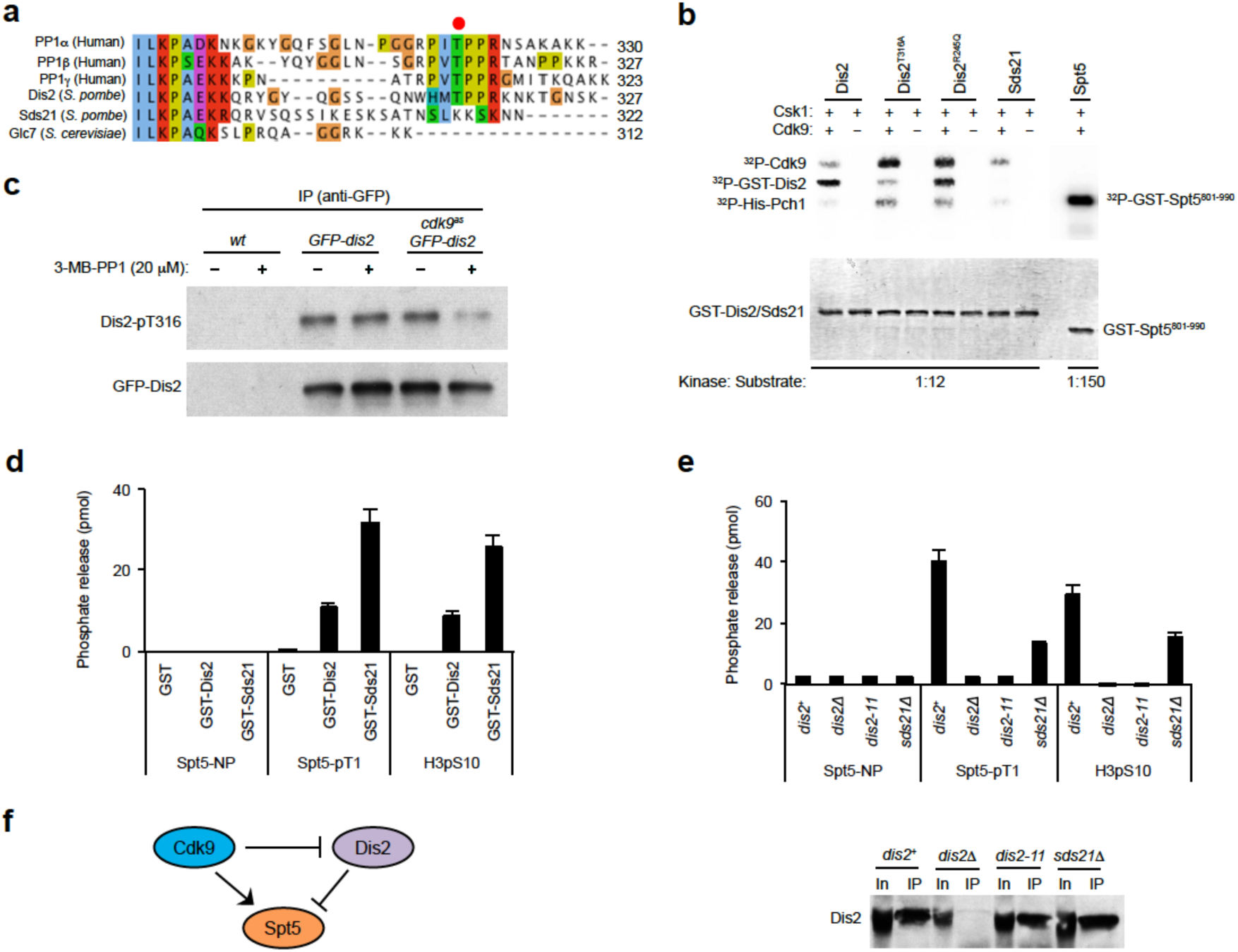
A Cdk9-Dis2-Spt5 circuit. **a**, Alignment of C-termini of human and fungal PP1 isoforms. In chemical-genetic screens, Thr320 in PP1α was identified as a potential target of Cdk1, and analogous residues in PP1β and PP1γ were identified as potential targets of Cdk9. This site of phospho-regulation (indicated by a red dot) is conserved in one of the two PP1 isoforms in fission yeast (Dis2-T316), but not in the other (Sds21), or in the budding yeast PP1 catalytic subunit Glc7. **b**, Phosphorylation of Dis2-T316 by Cdk9 *in vitro*. Purified, insect-cell derived Cdk9/Pch1 complexes were incubated at indicated molar ratios with purified, bacterially expressed GST-PP1 or GST-Spt5^801-990^ (containing the CTD), after activation by the CDK-activating kinase Csk1 (incubated alone in indicated lanes). In addition to wild-type Dis2, we tested Dis2^T316A^ and Dis2^R245Q^, the variant encoded by *dis2-11*. Autophosphorylation occurs on both Cdk9 and Pch1. (*Top*: autoradiogram; *bottom:* Coomassie-stained gel to confirm equal loading.) **c**, Cdk9-dependence of Dis2-T316 phosphorylation *in vivo*. Cells of *cdk9*^*as*^ strains, with or without GFP-tagged Dis2 expressed from the chromosomal *dis2*^+^ locus, were treated for 10 min with 20 μM 3-MB-PP1 or mock-treated, as indicated. Chromatin extracts were immunoprecipitated with anti-GFP antibodies and probed with antibodies specific for Dis2 phosphorylated at Thr316 (Dis2-T316P) or GFP. **d**, Spt5 dephosphorylation by purified PP1 *in vitro*. Purified GST-Dis2 and GST-Sds21 were incubated with a control phosphopeptide derived from histone H3 (“H3pS10”), an Spt5 CTD consensus phosphopeptide (“Spt5-pT1”), or a non-phosphorylated peptide of the same sequence (“Spt5-NP”). **e**, Spt5 dephosphorylation by PP1 isolated from fission yeast. A polyclonal anti-Dis2 antibody immunoprecipitates pSpt5 phosphatase activity from extracts of *dis2*^+^ but not *dis2* mutant cells. Note: the antibody cross-reacts with Sds21 in immunoblots but does not efficiently immunoprecipitate Sds21. For **d**, **e**, *n* = 3 biological replicates; error bars show standard deviation (s.d.). **f**, A Cdk9-Dis2-Spt5 circuit diagram.

The previously identified target of fission yeast Cdk9 is Thr1 in the CTD nonapeptide repeat T_1_P_2_A_3_W_4_N_5_S_6_G_7_S_8_K_9_ of Spt5^12,13^. A phosphopeptide containing this sequence was dephosphorylated by PP1s purified from bacteria (Fig. 1d) or isolated from S. *pombe* (Fig. 1e, Extended Data Fig. 1). We recovered similar amounts of Dis2 by immunoprecipitation from *dis2*^+^ and *dis2-11* cold-sensitive mutant cell extracts, but detected activity only in the former, consistent with the previous observation that the enzyme encoded by *dis2-11* has diminished activity even at permissive temperatures^10^. Finally, the amount of Dis2 we recovered from *sds21Δ* cells was similar to that from wild-type or *dis2-11* cells, but its activity was reduced, possibly suggesting a contribution by Sds21 to Dis2 activation. Together, these results indicate that Dis2 is a target of negative regulation by Cdk9 and a potential Spt5 phosphatase—an arrangement that predicts switch-like responses of pSpt5 to fluctuations in Cdk9 activity *in vivo* (Fig. 1f).

Consistent with this prediction, pSpt5 was rapidly lost after 3-MB-PP1 addition to exponentially growing *cdk9*^*as*^ cells (T_1/2_ ≈ 20 sec) (Fig. 2a, and accompanying paper by Booth *et al*.). A comparison of Spt5 dephosphorylation kinetics after Cdk9 inhibition in *dis2*^+^ versus *dis2-11* cells revealed an ~2-fold decrease in the dephosphorylation rate due to the *dis2-11* mutation at a permissive temperature of 30°C, and an ~4-fold decrease in *cdk9*^*as*^ *dis2-11* cells shifted to 18°C prior to 3-MB-PP1 addition (Fig. 2a, Extended Data Fig. 2a). Rapid decay at 18°C was restored by ectopic expression of active Dis2 in *dis2-11* cells (Fig. 2b). These results indicate that pSpt5 turnover is dependent on Dis2 activity in vivo.

**Figure 2.**
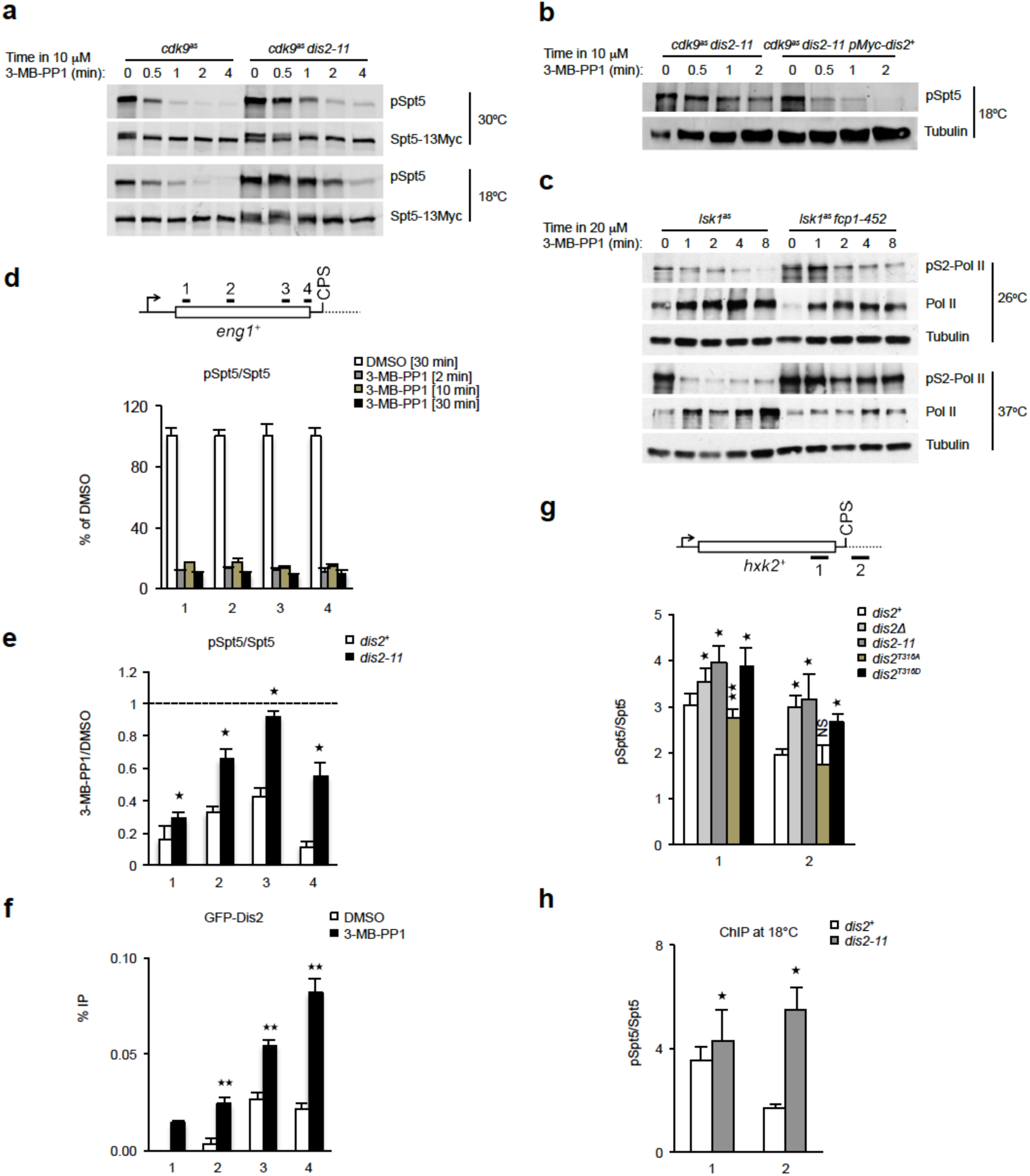
Cdk9 and Dis2 regulate Spt5 phosphorylation *in vivo*. **a**, Dis2 inactivation stabilizes pSpt5 after Cdk9 inhibition. Fission yeast strains—*cdk9*^*as*^ *spt5-13Myc dis2*^+^ or *cdk9*^*as*^ *spt5-13Myc dis2-11*—were grown to mid-log phase at 30°C and shifted to 18°C (bottom) or not shifted (top) for 10 min, before addition of 10 μM 3-MB-PP1, after which cultures were sampled at indicated times and subjected to immunoblot analysis with anti-pSpt5 or anti-Myc antibodies. **b**, Ectopic expression of wild-type Dis2 restores rapid Spt5 dephosphorylation kinetics in a *dis2* mutant. Anti-pSpt5 immunoblots from *cdk9*^*as*^ *dis2-11* cells shifted to 18°C and treated with 10 μM 3-MB-PP1 for indicated times, without or with expression of Myc-Dis2 from a plasmid. **c**, Fcp1 inactivation stabilizes Rpb1 Ser2 phosphorylation after Lsk1 inhibition. Fission yeast strains—*lsk1*^*as*^ or *lsk1*^*as*^ *fcp1-452*—were shifted to 37°C (or not shifted), treated for the indicated time with 20 μM 3-MB-PP1, and analyzed by immunoblotting for Rpb1-Ser2 phosphorylation. Note: CTD dephosphorylation leads to increased reactivity with the 8WG16 antibody used to detect total Pol II. **d**, Rapid pSpt5 turnover on chromatin. ChIP-qPCR analysis of pSpt5 versus total Spt5 crosslinking at the *eng1*^+^ gene after 3-MB-PP1 treatment for various times (expressed as a percentage of signal in the absence of inhibitor). **e**, Dis2 inactivation stabilizes chromatin-associated pSpt5. Either *cdk9*^*as*^ *spt5-13myc* or *cdk9*^*as*^ *spt5-13myc dis2-11* cells were shifted to 18°C and treated with 10 μM 3-MB-PP1 or mock-treated with DMSO for 2 min and subjected to ChIP-qPCR analysis at *eng1*^+^ for pSpt5 and total Spt5 (anti-Myc). The signal ratios between the two treatments were plotted for each condition. (Note: higher residual pSpt5 in *cdk9*^*as*^ *dis2*^+^ cells, compared to those analyzed in Fig. 2d, may reflect less efficient dephosphorylation at 18°C, relative to 30°C.) **f**, Suppression of Dis2 recruitment to transcribed chromatin by Cdk9. ChIP-qPCR analysis of GFP-Dis2 crosslinking at *eng1*^+^ in *cdk9*^*as*^ *GFP-dis2* cells treated for 10 min with 10 μM 3-MB-PP1. **g**, pSpt5 on chromatin in *dis2* mutants. ChIP-qPCR analysis of indicated strains upstream and downstream of CPS on *hxk2*^+^ gene at 30°C. **h**, Comparison of pSpt5:Spt5 ratio upstream and downstream of CPS on *hxk2*^+^ gene in *dis2*^+^ and *dis2-11* cells at 18°C. For **d-h**, *n* = 4 biological replicates; error bars show standard deviation (s.d.); asterisks indicate p-values (Student’s t-test: [*] *p* < 0.05; [**] *p* < 0.001; [NS] not significant) between wild-type (*dis2*^+^) and mutant (*dis2Δ, dis2-11, dis2*^*T316A*^ or *dis2*^*T316D*^) cells (**e**, **g**, **h**), or between 3-MB-PP1 and DMSO treatment (**f**).

Inhibition of the Cdk12 orthologue Lsk1 by 3-MB-PP1 treatment of *lsk1*^*as*^ cells^11^ had no effect on pSpt5 (Extended Data Fig. 2b) but rapidly diminished phosphorylation on Ser2 (pS2) of the Pol II CTD heptad repeat Y_1_S_2_P_3_T_4_S_5_P_6_S_7_. This mark became refractory to Lsk1 inhibition at 37°C in cells that harbored a temperature-sensitive mutation in *fcp1*^14^, which encodes a conserved pS2-specific phosphatase^15,16^ (Fig. 2c). The rate of pS2 decay in *lsk1*^*as*^ cells was unaffected by the *dis2-11* mutation (Extended Data Fig. 2c) and, conversely, Fcp1 inactivation had no effect on pSpt5 stability in *cdk9*^*as*^ strains (Extended Data Fig. 2d). Therefore, orthogonal CDK-phosphatase pairs govern pS2 and pSpt5, possibly to allow independent regulation of the two modifications, both of which are implicated in transcription elongation^17^.

Dis2 also influences pSpt5 turnover on chromatin; by ChIP-quantitative PCR (ChIP-qPCR) analysis, pSpt5 became nearly undetectable on *eng1*^+^ and *aro1*^+^ genes within 2 min of 3-MB-PP1 addition to *cdk9*^*as*^ *dis2*^+^ cells (Fig. 2d, Extended Data Fig. 3a, b), but persisted in *cdk9*^*as*^ *dis2-11* cells at 18°C (Fig. 2e, Extended Data Fig. 3c, d). Cdk9 inhibition stimulated GFP-Dis2 recruitment to regions where pSpt5 was stabilized by Dis2 inactivation (Fig. 2f). Dis2 chromatin association was also enhanced by *cdk9ΔC*, which removes a carboxy-terminal region of Cdk9 needed for its efficient recruitment to chromatin^18^, and by *cdk9*^*T212A*^, which prevents CdkQ-activating phosphorylation^19^ (Extended Data Fig. 4a). Therefore, CdkQ restricts Dis2 recruitment to chromatin in addition to its ability to inhibit Dis2 activity; both functions would promote switch-like changes in pSpt5 levels on genes in response to changes in Cdk9 activity.

**Figure 3.**
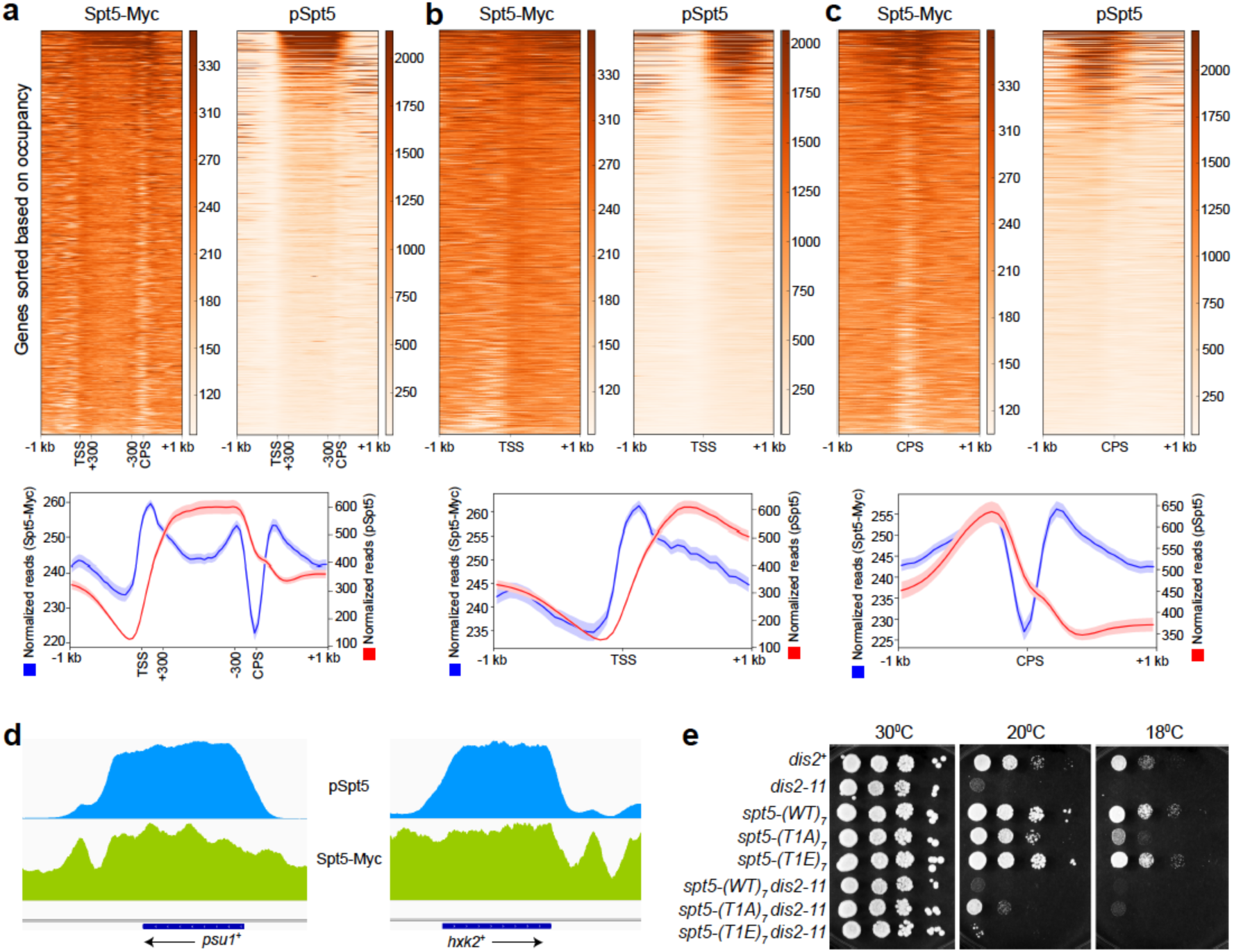
The Spt5 CTD is phosphorylated in gene bodies and unphosphorylated in the termination zone. **a**, pSpt5 distribution on transcribed genes. Heatmaps (top) and metagene analysis (bottom) reveal the pattern of Spt5-Myc and pSpt5 occupancy across transcribed genes, filtered to exclude those with neighboring genes closer than 1 kb. (Note: In this representation, the regions between +300 bp relative to the TSS and - 300 bp relative to the CPS have been scaled to allow comparisons among genes of different lengths.) **b**, pSpt5 distribution around the TSS. An unscaled, TSS-centered metagene analysis reveals a broad peak of pSpt5 downstream of TSS. **c**, Spt5 dephosphorylation in the termination zone. Unscaled metagene analysis centered at the CPS reveals a monotonic drop in pSpt5, but “twin peaks” of total, Myc-tagged Spt5. **d**, Spt5 dephosphorylation downstream of the CPS. Individual gene tracks show accumulation of Spt5-myc in the region downstream of the CPS with no concomitant peak of pSpt5. **e**, Suppression of *dis2-11* by an Spt5 CTD mutant that cannot be phosphorylated by Cdk9. Serial dilutions of indicated strains grown at 30°C, 20°C and 18°C. The *spt5* mutant alleles tested were: *spt5-(wt)*_*7*_, in which the CTD is truncated from 18 to seven wild-type, nonapeptide repeats; *spt5-(T1A)*_*7*_, in which each Thr1 position in the seven repeats is changed to alanine; and *spt5-(T1E)*_*7*_, in which each Thr1 is changed to glutamic acid.

**Figure 4.**
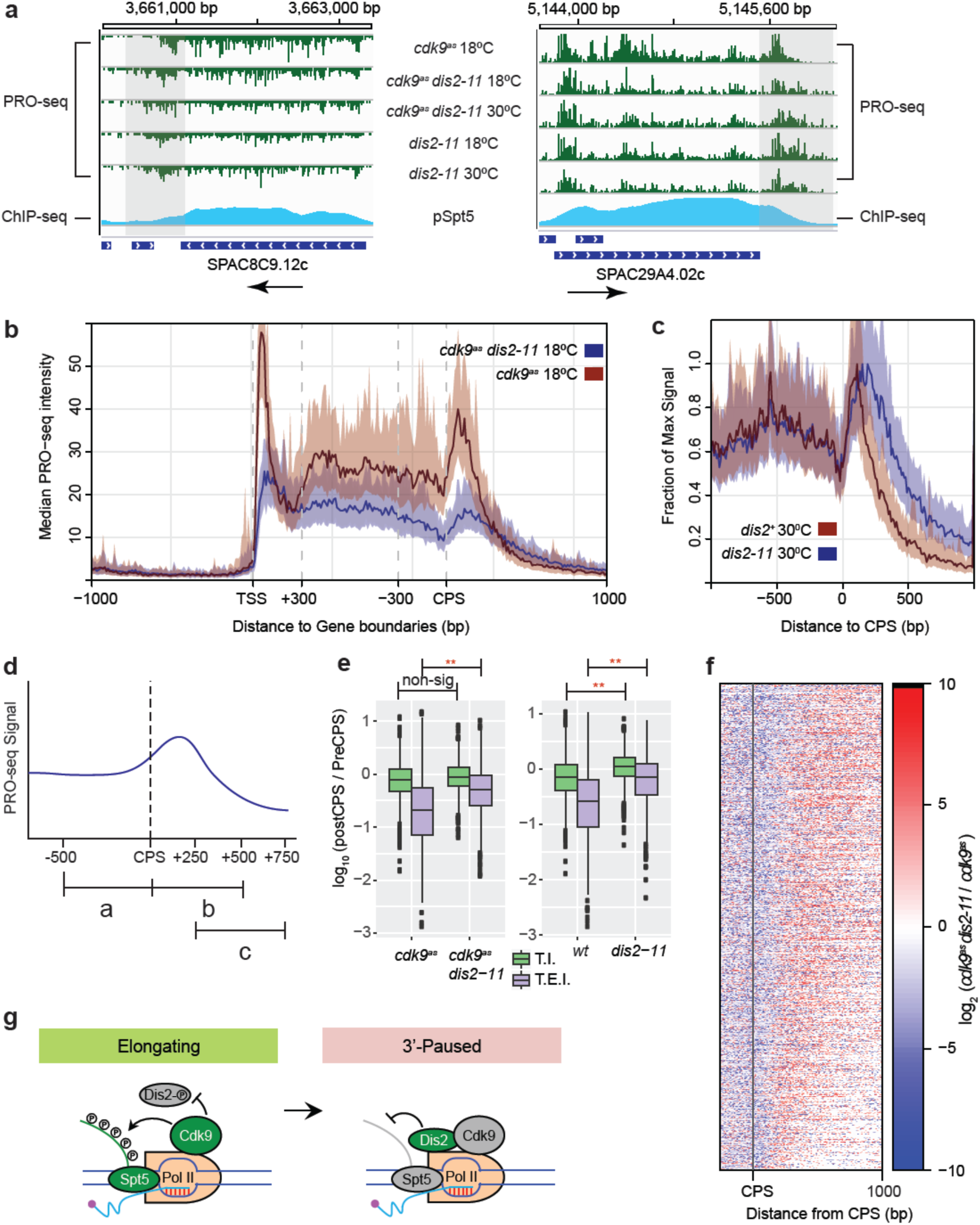
Loss of Dis2 function impairs termination. **a**, Transcription beyond normal termination zone in *dis2-11* cells under multiple conditions. Representative gene browser tracks where transcription terminates within a narrowly defined zone in *dis2*^+^ cells, but extends ~500 bp further downstream in *dis2-11* cells at both 18°C and 30°C. Alignment with ChIP-seq tracks reveals correlation between loss of pSpt5 and Dis2-dependent termination. **b**, Pleiotropic, genome-wide effects on Pol II dynamics due to *dis2-11* mutation. Metagene analysis of PRO-seq read distributions in *cdk9*^*as*^ *dis2*^+^ and *cdk9*^*as*^ *dis2-11* strains reveals multiple differences in *dis2-11*, relative to *dis2*^+^ cells. These analyses were performed in the absence of 3-MB-PP1 treatment; under this condition, PRO-seq read distributions were not significantly different between *cdk9*^+^ and *cdk9*^*as*^ cells, as shown in the accompanying paper by Booth *et al*. Note: To compare genes of different lengths, regions between +300 bp relative to TSS and ‐300 bp relative to CPS are scaled. **c**, CPS-centered metagene analysis comparing PRO-seq read distributions in wild-type and *dis2-11* cells reveals relative increase in transcription downstream of the CPS in the mutant at 30°C. Peak heights (y axis) were scaled as a fraction of maximum signal in each condition, whereas position along the gene (*x* axis) was unscaled. **d**, Loss of Dis2 function affects two metrics of termination efficiency, shown schematically: Termination Index (T.I.), the ratio of signals in the regions 500 bp downstream and upstream of the CPS (b/a); and Termination Elongation Index (T.E.I.), the ratio of signals in the region between +250 bp and +750 bp relative to CPS to that in the region 500 bp upstream of CPS (c/a). **e**, Box plots show that *dis2-11* mutation causes a significant increase in T.E.I. in both *cdk9*^*as*^ (left) and *cdk9*^+^ (WT, right) backgrounds, and in T.I. in *cdk9*^+^ cells (Student’s t-test, *p* < 0.01). **f**, Heatmaps showing change in PRO-seq read distribution due to *dis2-11* mutation. Genes were ranked by decreasing T.E.I. in *cdk9*^*as*^ *dis2-11* at 18°C, a measure of termination-window size. Dis2 impairment expands this zone relative to *cdk9*^*as*^ at 18°C. All genes in **b-f** were required to be active and at least 1 kb from nearest genes on the same strand to eliminate effects of nearby initiation and run-through transcription (n = 919). **g**, A transcription exit network comprising Cdk9, Dis2 and Spt5. At or near CPS, Dis2 becomes active due to a drop in Cdk9 activity and triggers Spt5 dephosphorylation, to facilitate 3’-pausing and termination.

ChIP-qPCR analysis of *hxk2*^+^ and *rps17a*^+^ genes, in *cdk9*^+^ cells with different *dis2* alleles at 30°C, revealed statistically significant increases in pSpt5 in the loss-of-function mutants *dis2Δ, dis2-11* and *dis2*^*T316D*^ (aspartic acid substitution to mimic constitutive Thr316 phosphorylation^4^), relative to *dis2*^+^ cells and cells harboring a CDK-refractory, *dis2*^*T316A*^ allele (Fig. 2g, Extended Data Fig. 4b). Stabilization of pSpt5 was more prominent in *dis2-11* cells shifted to 18°C, and occurred preferentially in regions downstream of the CPS (compare Fig. 2h to 2g, Extended Data Fig. 4c to 4b). The relative increases in chromatin-associated pSpt5 in *dis2-11* and *dis2Δ* cells at 30°C correlated with the degree of bulk pSpt5 stabilization after Cdk9 inactivation (Fig. 2a, Extended Data Fig. 5a). We suspect that *dis2-11* is more severely affected than *dis2Δ* because loss of Dis2 protein allows more effective compensation by other phosphatases such as Sds21. An *sds21Δ* mutation, however, did not delay pSpt5 decay, and Cdk9 inhibition did not increase chromatin recruitment of Sds21 (Extended Data Fig. 4d, 5b), indicating that, in wild-type cells, Dis2 is the major PP1 isoform that regulates pSpt5 in opposition to Cdk9.

In the accompanying paper, PRO-seq analysis uncovered a rate-limiting role for Cdk9 in Pol II elongation (Booth *et al.)*. That function is likely to depend on Spt5, depletion of which slowed elongation and caused Pol II accumulation in upstream gene regions in fission yeast^20^, and disrupted coupling between 3’-processing and termination in budding yeast^21^. In ChIP-seq analysis, we detected total Spt5 and pSpt5 in the bodies of Pol II-transcribed genes (Fig. 3a), with the phosphorylated form accumulating further downstream of the transcription start site (TSS) (Fig. 3b). The patterns diverged again downstream of the CPS; a 3’ peak of apparently unphosphorylated Spt5 is prominent in metagene profiles (Fig. 3c) and individual gene tracks (Fig. 3d), and correlates with a peak of paused, Ser2-phosphorylated Pol II detected by Winston and co-workers^20^ (Extended Data Fig. 6a-f).

The Spt5-Myc metagene profile features a V-shaped depression centered just upstream of the CPS (Fig. 3c), which is also seen in the Pol II pattern derived from published ChIP-seq data^20^ (Extended Data Fig. 6c), and in a ChIP-seq analysis of Spt5 in budding yeast, where it corresponds to a peak of Spt5 cross-linking to the nascent transcript^21^. Although exchange of phosphorylated for unphosphorylated Spt5 is possible, Spt5’s tight association with the Pol II clamp^22^ favors active dephosphorylation as a more likely explanation for the divergence between pSpt5 and Spt5-Myc occupancy downstream of the CPS—the same region where pSpt5 was preferentially stabilized by Dis2 inactivation. Moreover, Spt5 and Dis2 interact genetically; replacement of Thr1 with alanine in a truncated, seven-repeat Spt5 CTD—*spt5-(T1A)*_*7*_, which by itself imparts cold-sensitivity^23^—partially suppressed cold-sensitive lethality due to *dis2-11*, whereas a phosphomimetic *spt5-(T1E)*_*7*_ mutation (which did not affect growth on its own^23^) exacerbated this phenotype (Fig. 3e, Extended Data Fig. 7).

We next performed PRO-seq analysis^6^ to uncover effects of Dis2 inactivation on the distribution of transcriptionally engaged Pol II (Fig. 4, Extended Data Fig. 8). On individual genes, transcribing Pol II decreased dramatically within a narrowly defined zone following the CPS in *dis2*^+^ cells, but this zone extended ~500 bp further downstream in *dis2-11* cells at both 18°C and 30°C. Alignment of PRO-seq and ChIP-seq read distributions suggested correlation between the zone of Dis2-dependent termination and Spt5 dephosphorylation (Fig. 4a). Metagene analysis of PRO-seq data (Fig. 4b, c, Extended Data Fig. 9a, b) revealed pleiotropic effects of the *dis2-11* mutation: 1) attenuated pausing in promoter-proximal regions^5^; 2) decreased density of transcribing Pol II throughout gene bodies; and 3) increased transcription beyond the CPS, both in absolute terms and relative to transcription of upstream regions. All three defects were apparent at both 18°C and 30°C, indicating that the effects of Dis2 inactivation on Pol II distribution are constitutive. Loss of viability occurs only at low temperatures, however, and is likely due in part to inappropriate persistence of pSpt5 (Fig. 2a), as suggested by the partial rescue achieved with the *spt5-(T1A)*_*7*_ mutation (Fig. 3e).

To quantify termination defects due to Dis2 impairment, we defined two metrics based on PRO-seq read distribution: Termination Index (T.I.), the ratio of signals in the regions 500 bp downstream and upstream of the CPS; and Termination Elongation Index (T.E.I.), the signal ratio in the regions 250-750 bp downstream and 500 bp upstream of the CPS (Fig. 4d). The *dis2-11* mutation caused statistically significant increases in T.I. in *cdk9*^+^ cells, and in T.E.I. in both *cdk9*^+^ and *cdk9*^*as*^ backgrounds (Fig. 4e). A heatmap analysis of genes ranked by T.E.I. revealed termination-zone expansion consistent with decreased termination efficiency upon loss of Dis2 function (Fig. 4f, Extended Data Fig. 9c). In the accompanying paper, Booth *et al*. show that Cdk9 inhibition has the opposite effect—decreasing both T.I. and T.E.I.—consistent with functional antagonism between the kinase and phosphatase.

Recent studies suggest an ordered series of events at 3’-ends of protein-coding genes: 1) slowing of elongation, 2) increased pS2 as a consequence of this pause, 3) cleavage and polyadenylation factor (CPF) recruitment to the pS2-containing CTD, 4) CPF-dependent cleavage and 5) termination facilitated by the 5’→3’ exoribonuclease XRN2/Rat1^12,3,21,24,25^ This sequence can be initiated ectopically by a block to transcription imposed *in cis*^*2*^, and pS2 is elevated in 5’ gene regions by mutations that decrease the intrinsic elongation rate of Pol II^26^, but a physiologic trigger remains unknown. Both Spt5 and PP1 are implicated in this transition—the former as a regulator of Pol II processivity and rate^20,21,27^, the latter as a CPF component^28,29^. We now show that Spt5 and the PP1 isoform Dis2 are both substrates of Cdk9, a positive regulator of elongation^30^. The enzyme-substrate relationships we define among Cdk9, Dis2 and Spt5 recapitulate an important cell-cycle regulatory module^4^, and suggest a model of “transcriptional exit” (Fig. 4g): Dis2-dependent Spt5 dephosphorylation upon reversal of a Cdk9-PP1 switch, leading to slowing of Pol II to facilitate its capture and dissociation by XRN2/Rat1.

## Acknowledgements

We thank I.M. Hagan, B. Schwer, S. Shuman, M. Yanagida and M.J. O’Connell for providing yeast strains and antibodies; K. M. Shokat for providing 3-MB-PP1; C. Zhang for guidance in AS-allele optimization; and N. Steinbach and R. Parsons for advice and assistance in phosphatase activity measurements. J.C.T. was supported by Canadian Institutes of Health Research grant MOP-130362 and by a fellowship from Fond de recherche Quebec Santé (3315). This work was supported by National Institutes of Health grants GM25232 to G.T.B. and J.T.L., and GM104291 to R.P.F.

## Author Contributions

P.P., G.T.B. and M.S. designed and performed experiments and performed data analysis. B.B. and J.C.T. performed data analysis. P.P., G.T.B., J.T.L. and R.P.F. prepared the manuscript.

## Author Information

Correspondence and requests for materials should be addressed to

## METHODS

### Yeast strains and standard methods

Fission yeast strains used in this study are listed in Extended Data Table 1. New strains were generated by standard techniques^31^. Cells were grown in YES medium at 30°C unless otherwise specified.

### Immunological methods

Antibodies used in this study recognized pSpt5 or total Spt5^13^, Dis2-pT316^4^ or total Dis2^32^, Myc epitope (EMD Millipore, 05-724), total Pol II (BioLegend, MMS-126R), Pol II pSer2 (Abcam, ab5095), α-tubulin (Sigma, T-5168) and green fluorescent protein (GFP) (Invitrogen, rabbit polyclonal (A11122) or Santa Cruz, mouse monoclonal (sc-9996)). Proteins were visualized in immunoblot analysis either by enhanced chemiluminescence (ECL, HyGLO HRP detection kit, Denville Scientific, E250) or with Odyssey Imaging System (LI-COR Biosciences).

### Kinase and phosphatase assays

Kinase assays were performed with purified proteins (Cdk9/Pch1 complex and GST-Spt5^801-990^) and [γ-^32^P] ATP (PerkinElmer, BLU002A250UC), as described previously^19^. GST-PP1s were expressed in *E. coli* at 16°C for 16 h and purified with Glutathione Sepharose 4 Fast Flow beads^33^. To measure protein phosphatase activity, PP1 isoforms (GST-Dis2 or GST-Sds21) purified from *E. coli* (2 μg) or immunoprecipitated from fission yeast extracts (4 mg total protein) were incubated with 50 μM peptide (Spt5-NP, Spt5-pT1 or H3pS10) at 37°C for 1 h. Colorimetric assays were performed in triplicate using BioMOL Green (Enzo Life Sciences, BML-AK111) in 25 mM HEPES (pH 7.5), 100 mM NaCl_2_, 1 mM MnCfe and 1 mM DTT, in 96-well plates as described in manufacturer’s protocol.

### Chromatin immunoprecipitation analysis

Immunoprecipitation and chromatin immunoprecipitation (ChIP) were carried out by published methods^18,34,35^. Quantitative PCR was performed with USB VeriQuest SYBR Green qPCR Master Mix (2x) (Affymetrix, 75600) in 386-well plates. To measure the dependency of Dis2-Thr316 phosphorylation on Cdk9 *in vivo*, cells expressing GFP-Dis2 from the chromosomal *dis2* locus in either *cdk9*^+^ or *cdk9*^*as*^ background were grown at 30°C to a density of ~ 1.5 × 10^7^ ml^-1^, treated with either DMSO or 10 μM 3-MB-PP1 for 10 min, and crosslinked with 1% formaldehyde for 15 min at 25°C. Chromatin extracts were prepared according to a standard protocol for ChIP sample preparation^18,34,35^ and 5 mg total protein was subjected to immunoprecipitation with anti-GFP antibody (sc-9996) and immunoblot analysis with either anti-GFP or anti-Dis2-pT316 antibody^4^.

### Chemical genetics

To measure rates of Spt5 and Rpb1 dephosphorylation after CDK inhibition, cells were grown at 30°C in YES medium to a density of ~1.2 × 10^7^ cells/ml, harvested by centrifugation at 25°C, resuspended in fresh YES (preincubated at desired temperature) and incubated for 10 min before addition of DMSO or 3-MB-PP1. At intervals after addition of DMSO or 3-MB-PP1, cells (~0.6 × 10^8^) were transferred directly to tubes containing 500 μl 100% (w/v) trichloroacetic acid (TCA) and harvested by centrifugation. Protein extracts were prepared in the presence of 20% (w/v) TCA^36^ and processed for immunoblot analysis with appropriate antibodies.

### ChIP-seq

Multiplexed ChIP-seq libraries were prepared using the Illumina TruSeq DNA Sample Preparation kit v2 with 75 ng of input or IP DNA and barcode adaptors. Pairedend sequencing (50-nt reads) was performed on an Illumina HiSeq 2000 (Genome Quebec Innovation Centre, McGill University). After adaptor trimming and quality control, processed FASTQ files were aligned to the S. *pombe* genome using Bowtie2^37^ (Galaxy Version 2.2.6.2). Aligned sequences of each biological replicate were fed into MACS2^38^ (Galaxy Version 2.1.1.20160309.0) to call peaks from alignment results. Generated “bedgraph treatment” files were concatenated (Galaxy Version 1.0.1) to combine replicates of each sample, converted into bigwig using “Wig/BedGraph-to-bigWig converter” (Galaxy Version 1.1.0) and processed using computeMatrix (Galaxy Version 2.3.6.0) in DeepTools^39^ to prepare data for plotting a heatmap and/ or a profile of given regions. Heatmap and Metagene plots were generated using “plotHeatmap” (Galaxy Version 2.5.0.0) and “plotProfile” (Galaxy Version 2.5.0.0) tools, respectively.

### PRO-seq

For PRO-seq analysis, all samples, with the exception of the wild-type (*wt*) strain (prepared separately), were prepared simultaneously and in biological replicates. A detailed description of PRO-seq libraries can be found in the methods of the accompanying manuscript (Booth *et al*.). Briefly, biological replicates for each strain were derived from separately picked colonies. Cultures were grown in YES medium at 30°C overnight and diluted to an OD_600_ = 0.2. After reaching OD_600_ ≈ 0.5, an equal number of cells (based on OD) was set aside for all treatments (~10 ml culture). At this point, a fixed amount of thawed S. *cerevisiae* (50 μl, OD = 0.68) was spiked into each sample. Cells were spun down and resuspended in fresh YES medium, pre-conditioned at the desired temperature (30°C or 18°C) and allowed to incubate for 10 min. Samples were immediately spun down at 4 °C and subjected to permeabilization and library preparation as described by Booth *et al.*, using the standard 5’ and 3’ RNA adaptors^40^. Sequencing, alignment and processing of reads were conducted as described in the accompanying paper.

### Data Access

The raw and processed sequencing files have been submitted to the NCBI Gene Expression Omnibus (GEO; http://www.ncbi.nlm.nih.gov/geo/) (Accession number pending).

---

**Extended Data Figure 1.**
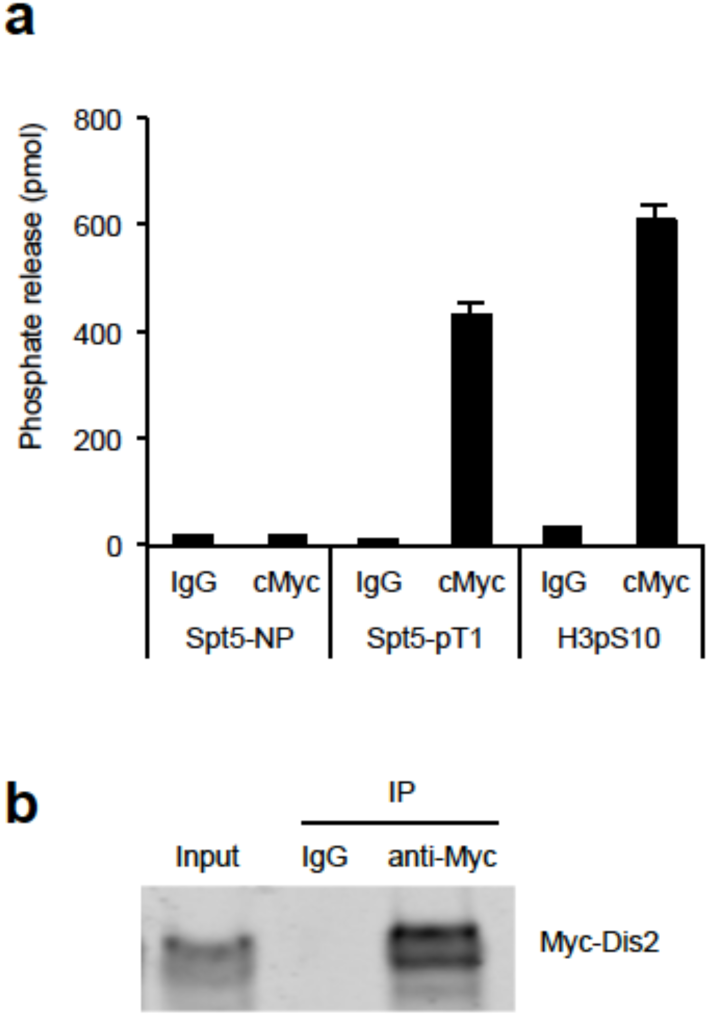
Dis2 expressed in fission yeast dephosphorylates Spt5-T1P *in vitro*. **a**, Anti-Myc immunoprecipitates from extracts of Myc-Dis2-expressing cells were tested for phosphatase activity towards the Spt5 CTD-derived phosphopeptide. *n* = 3 biological replicates; error bars show standard deviation (s.d.). **b**, Immunoblot to verify expression and immunoprecipitation of Myc-Dis2.

**Extended Data Figure 2.**
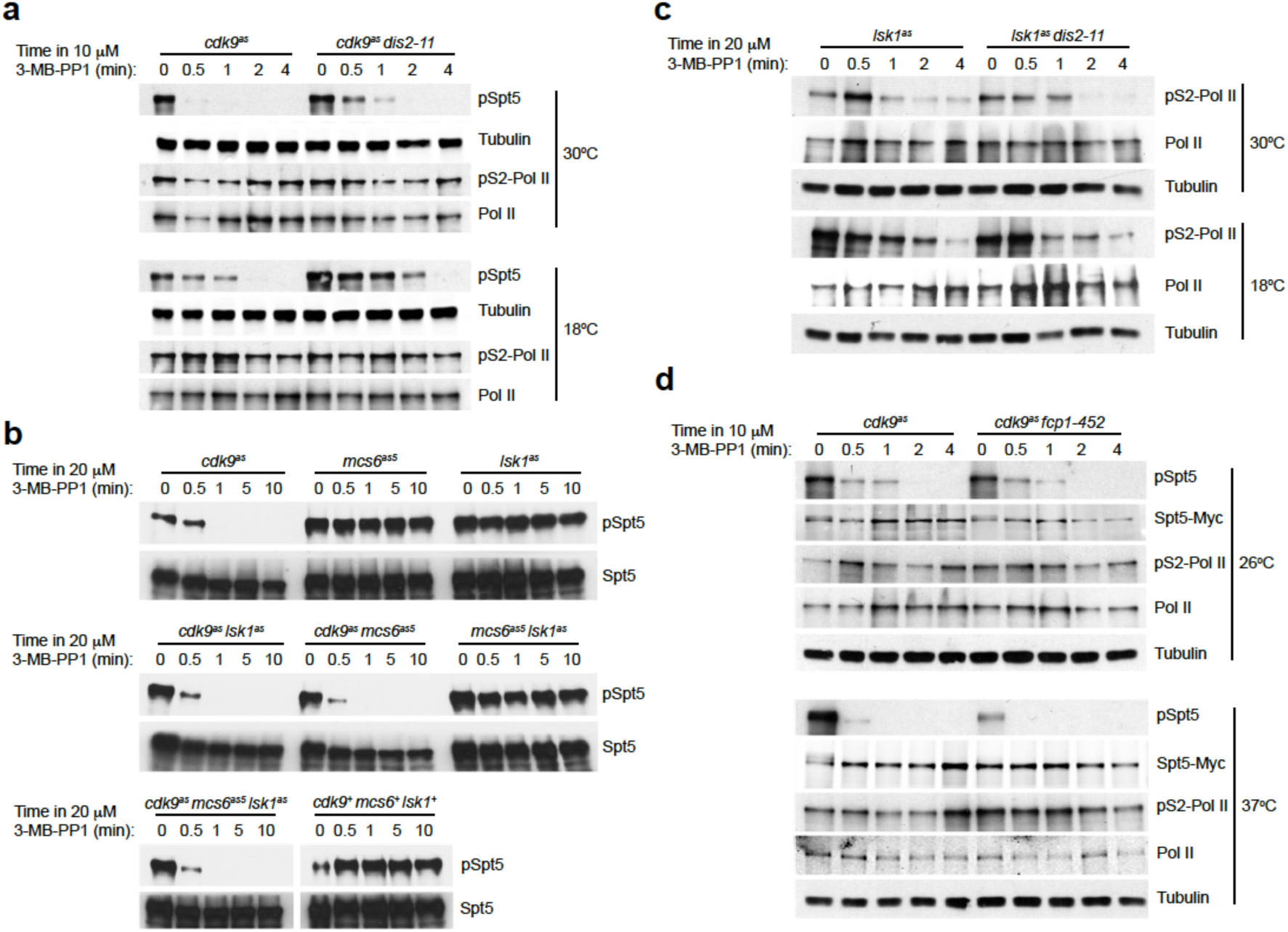
Distinct kinase-phosphatase circuits regulate phosphorylation of Spt5-Thr1 and Rpb1-Ser2 *in vivo*. **a**, Rapid kinetics of Spt5 dephosphorylation after Cdk9 inhibition, and stabilization of pSpt5 by Dis2 inactivation occur in *spt5*^+^ strain and do not depend on the C-terminal Myc-epitope tag. **b**, Spt5 dephosphorylation kinetics in single, pairwise and triple *cdk*^*as*^ mutants treated with 3-MB-PP1 indicate that Cdk9 is the sole kinase needed to phosphorylate this site *in vivo*. **c**, Dis2 activity is dispensable for Pol II CTD Ser2 dephosphorylation. As in Fig. 2a except that experiment was performed in *lsk1*^*as*^ cells, 20 μM 3-MB-PP1 was added, and extracts were probed for phosphorylation of the Ser2 position of the Rpb1 CTD (or tubulin as a loading control). **d**, Fcp1 activity is dispensable for Spt5 Thr1 dephosphorylation. As in Fig. 2c except that strains carried a *cdk9*^*as*^ allele and were tested for both pSpt5 (unaffected by Fcp1 inactivation) and pS2 (unaffected by Cdk9 inhibition).

**Extended Data Figure 3.**
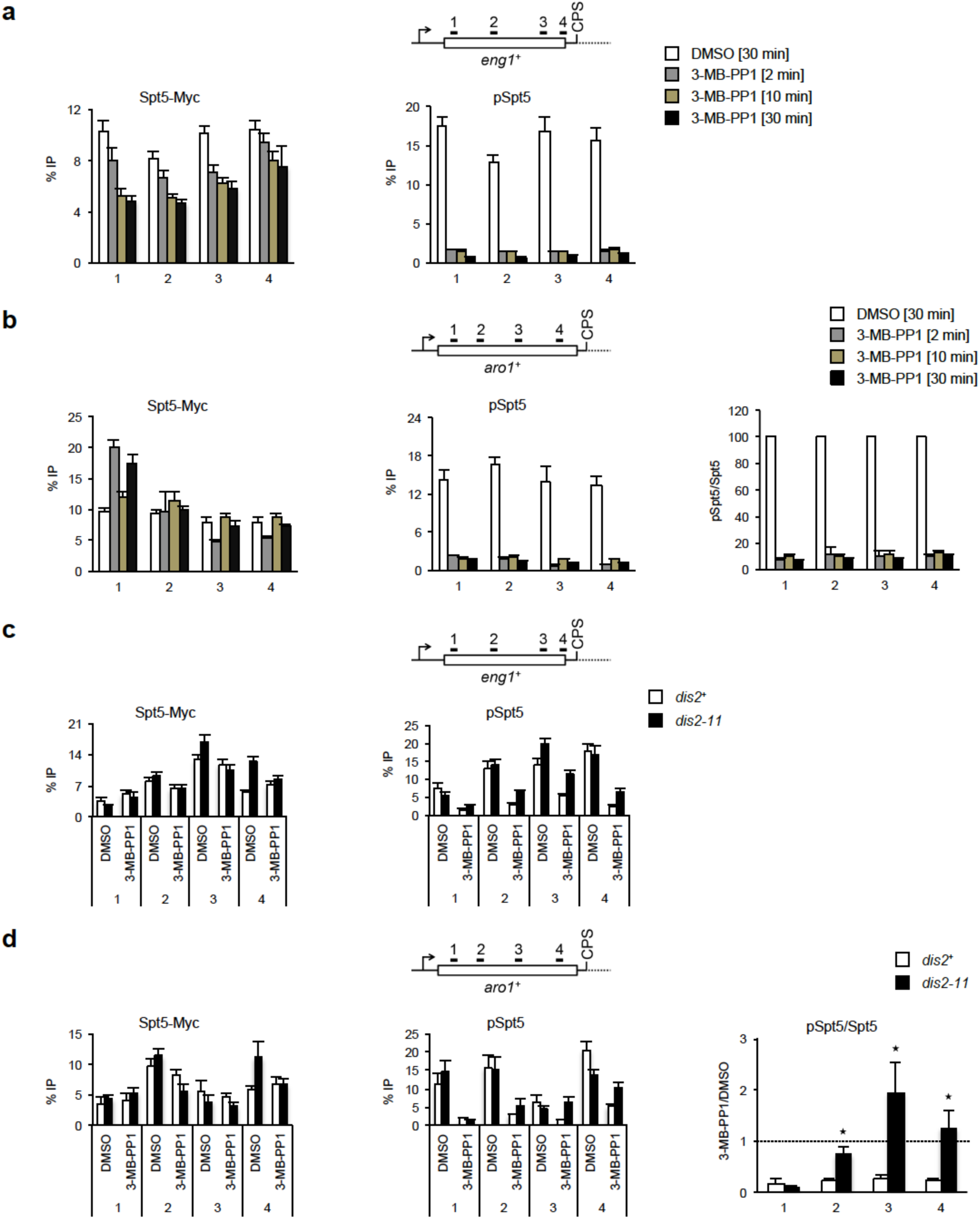
Rapid dephosphorylation of chromatin-associated Spt5 upon Cdk9 inhibition. **a**, Dephosphorylation of Spt5 on chromatin of *eng1*^+^ gene after Cdk9 inhibition (raw data for phospho‐ and total Spt5 from which ratios in Fig. 2d were calculated). **b**, ChIP-qPCR analysis of phosphoversus total Spt5 crosslinking at the *aro1*^+^ gene after 3-MB-PP1 treatment for various times. Left: absolute ChIP signals for anti-Myc; middle: absolute signals for anti-pSpt5; right: ratio of phospho-to total Spt5, expressed as a percentage of ratio in the absence of the inhibitor. **c**, Dis2 inactivation stabilizes pSpt5 on chromatin. Either *cdk9*^*as*^ *spt5-13Myc dis2*^+^ or *cdk9*^*as*^ *spt5-13Myc dis2-11* cells were shifted to 18°C and treated with 10 μM 3-MB-PP1 or mock-treated with DMSO for 2 min and subjected to ChIP-qPCR analysis at the *eng1*^+^ locus for pSpt5 and Spt5-Myc (raw data for phospho‐ and total Spt5 from which ratios in Fig. 2e were calculated). **d**, Same as (c) at *aro1*^+^ gene: ChIP-qPCR analysis of pSpt5 (left) and Spt5-Myc (middle) and the signal ratios in 3-MB-PP1versus DMSO-treated samples (right). For **a-d**, *n* = 4 biological replicates; error bars show standard deviation (s.d.).

**Extended Data Figure 4.**
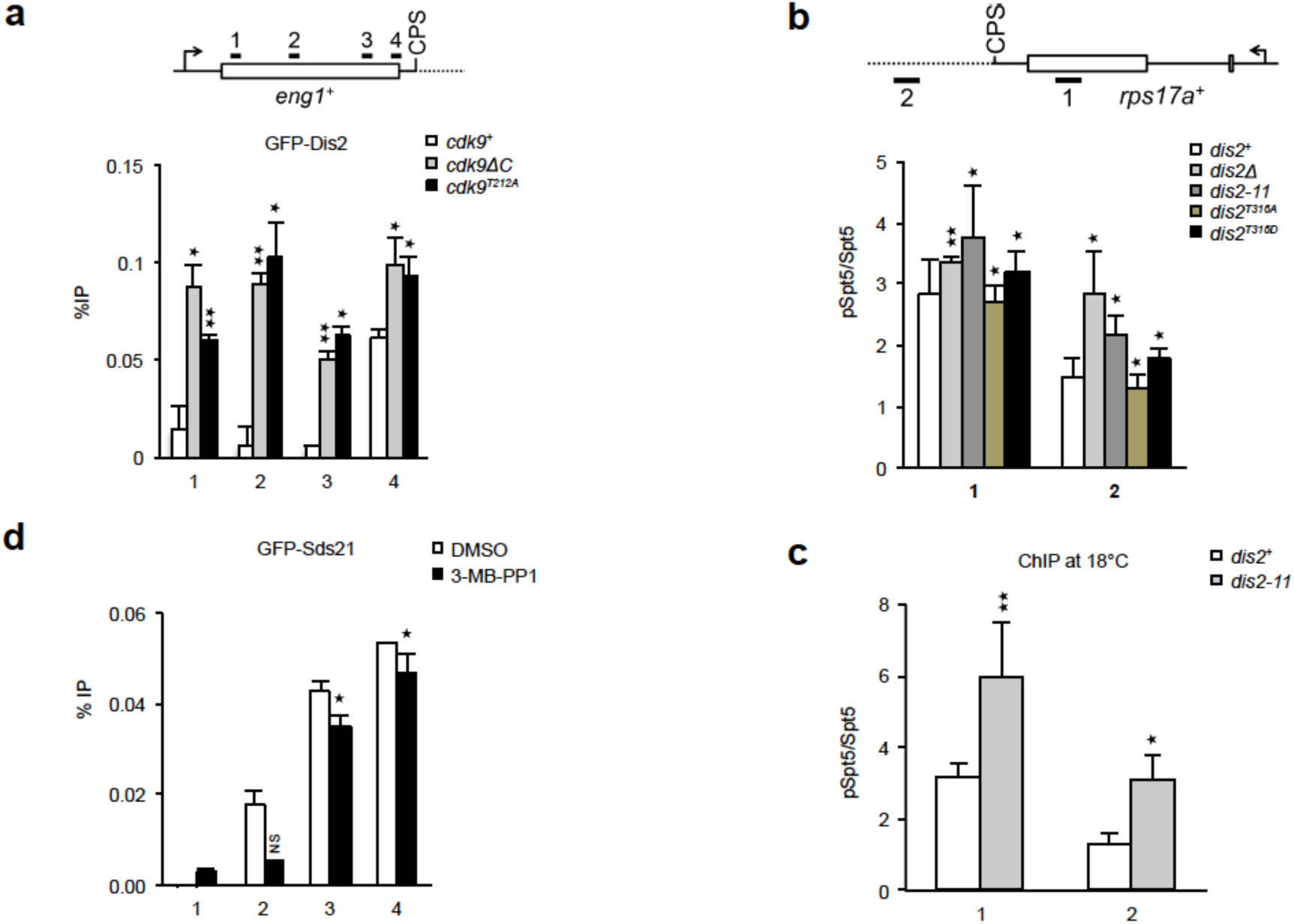
Cdk9 regulates PP1 recruitment to chromatin in isoform-specific fashion. **a**, Constitutive *cdk9* loss-of-function mutations increase GFP-Dis2 recruitment to chromatin. Dis2 occupancy at *eng1*^+^ locus analyzed in *cdk9*^+^ (“wt”) cells, a *cdk9ΔC* mutant and a *cdk9*^*T212A*^ mutant. **b**, Spt5 phosphorylation on chromatin in *dis2* mutants. ChIP-qPCR analysis of indicated strains upstream and downstream of CPS on *rps17a*^+^ gene at 30°C. **c**, Comparison of pSpt5:Spt5 ratio upstream and downstream of CPS on *rps17a*^+^ gene in *dis2*^+^ and *dis2-11* cells at 18°C. d, Anti-GFP ChIP at *eng1*^+^ in a *cdk9*^*as*^ *GFP-sds21* strain treated for 10 min with 10 μM 3-MB-PP1 reveals unchanged or slightly decreased Sds21 occupancy when Cdk9 is inhibited. For **a-d**, *n* = 4 biological replicates; error bars show standard deviation (s.d.); asterisks indicate p-values (Student’s t-test: [*] *p* < 0.05; [**] *p* < 0.001; [NS] not significant) (a) between wild-type (*cdk9*^+^) and mutant (*cdk9ΔC* or *cdk9*^*T212A*^), (b, c) between wild-type (*dis2*^+^) and mutant (*dis2Δ*, *dis2-11*, *dis2*^*T316A*^ or *dis2*^*T316D*^), or (d) between 3-MB-PP1 and DMSO.

**Extended Data Figure 5.**
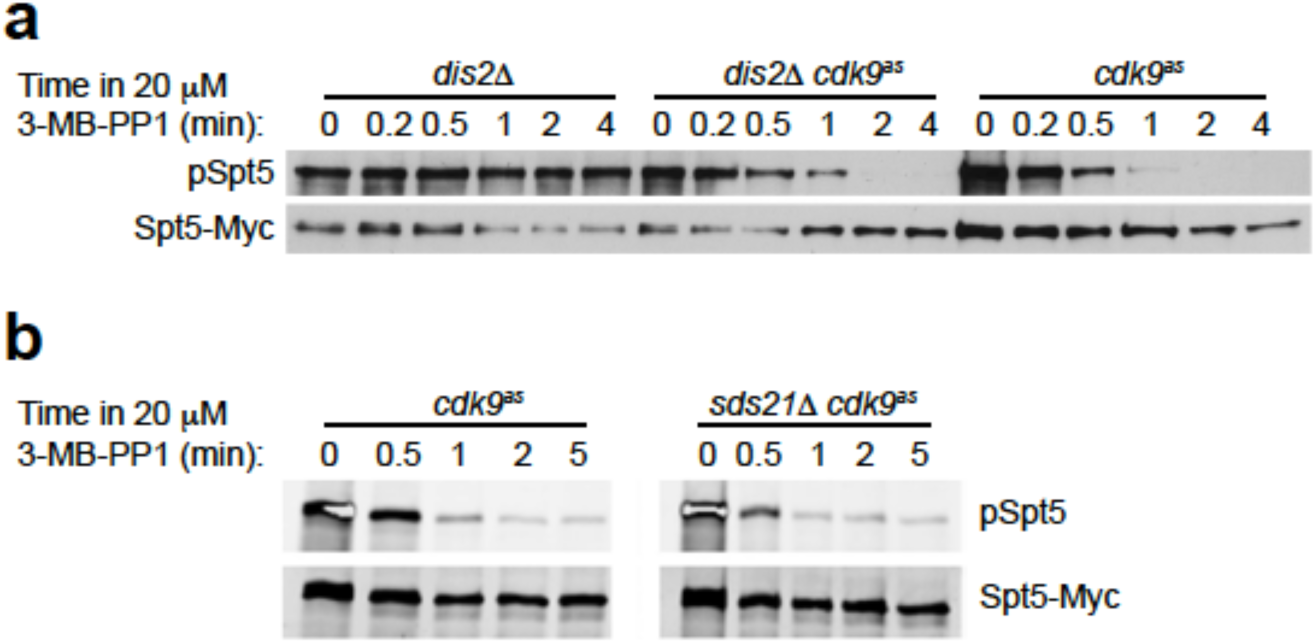
Spt5 dephosporylation kinetics in cells with a single PP1 isoform. **a**, Dephosphorylation of Spt5 after Cdk9 inhibition is retarded in *dis2Δ* strain, relative to a *dis2*^+^ strain. **b**, Spt5 dephosphorylation kinetics after Cdk9 inhibition are unaffected by *sds21* deletion in a *dis2*^+^ strain.

**Extended Data Figure 6.**
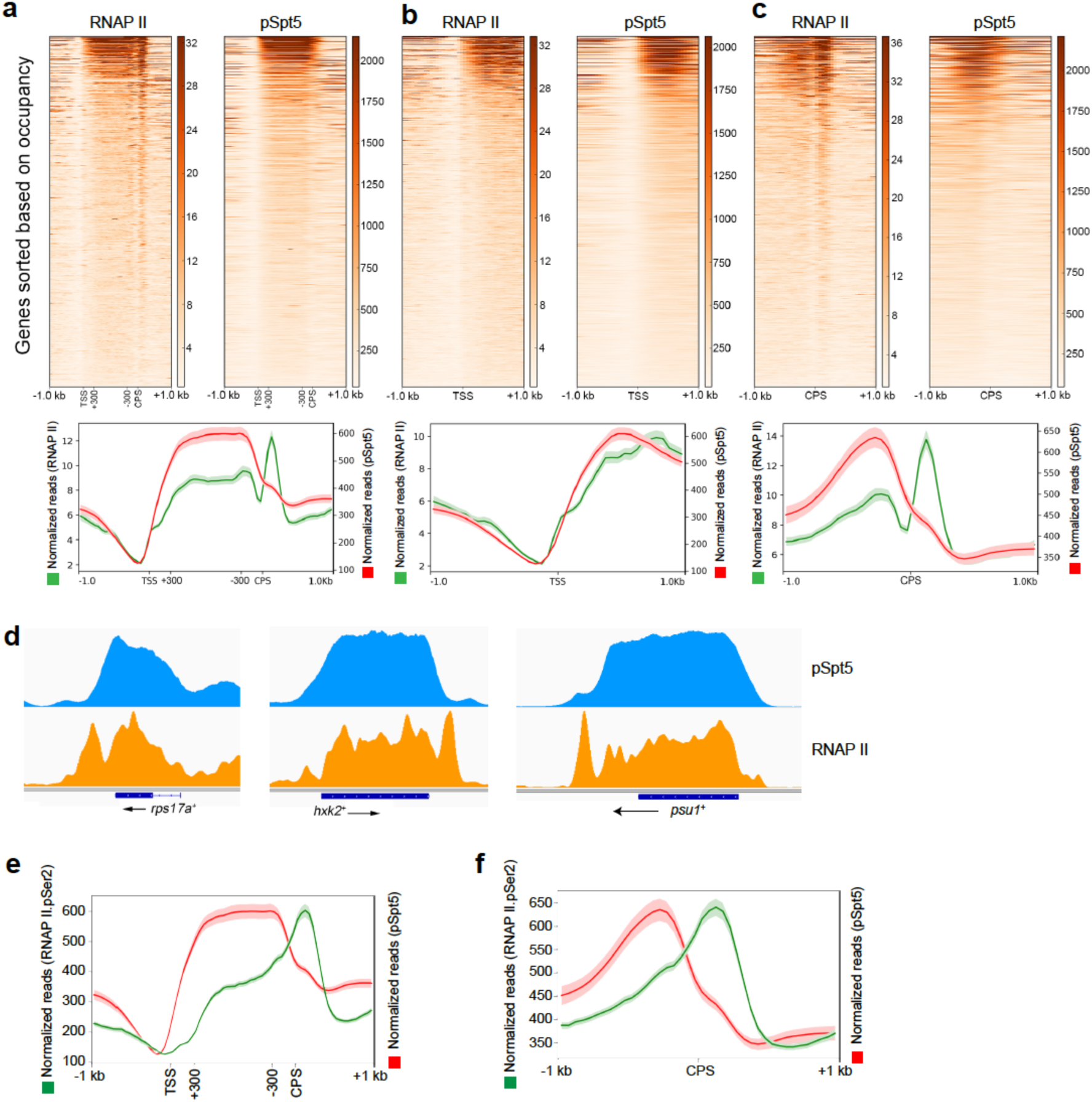
Pol II and pSpt5 distribution on chromatin. **a-c**, Heatmap (top) and metagene (bottom) analyses, scaled as in Fig. 3a-c, of Pol II ChIP-seq data compared with pSpt5 over entire gene (**a**), and in TSS (**b**) and CPS (**c**) regions. **d**, Genome browser tracks of representative genes, showing occupancy of pSpt5 and Pol II. **e**, Metagene analysis comparing distribution of pSpt5 and pS2 over entire gene, scaled as in (**a**). **f**, Metagene analysis comparing distribution of pSpt5 and pS2 centered around CPS, without scaling, as in (**c**).

**Extended Data Figure 7.**
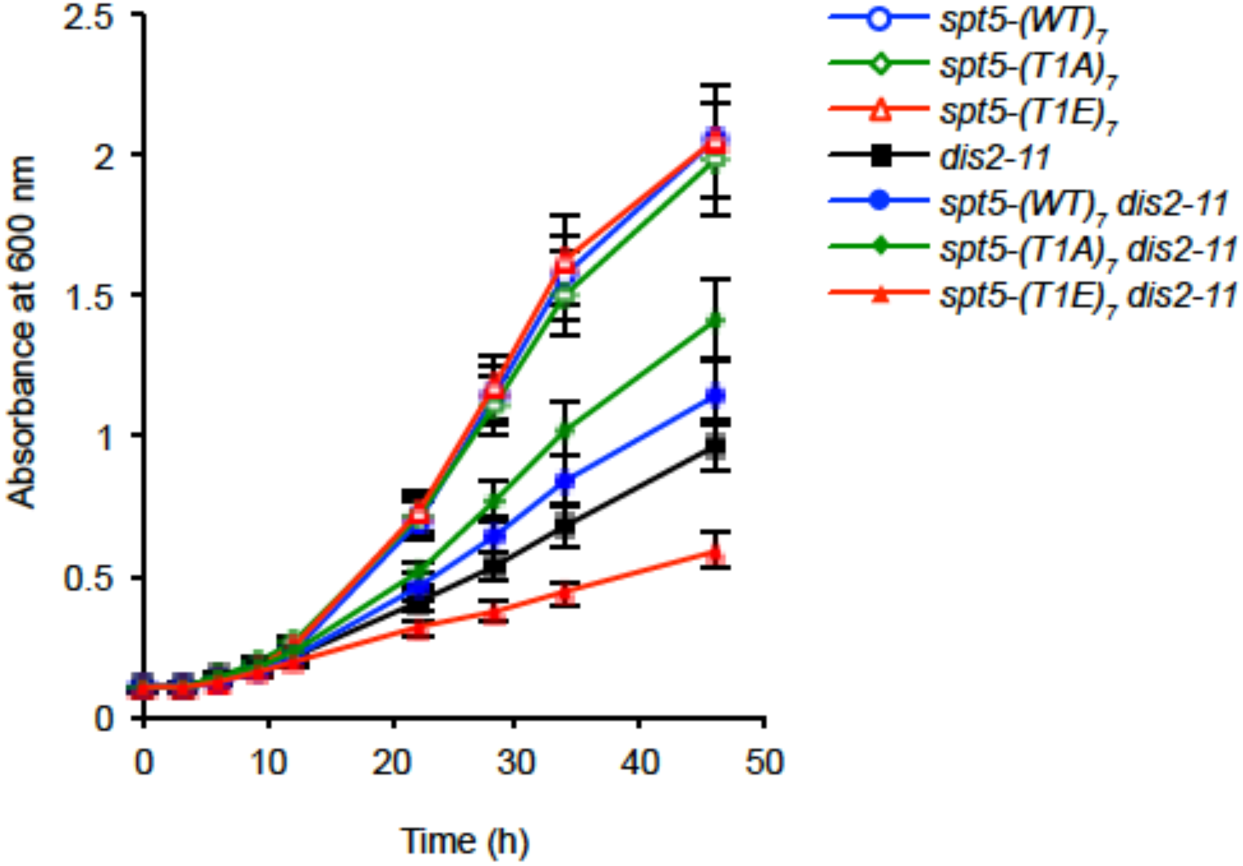
A non-phosphorylatable Spt5 suppresses conditional lethality of *dis2-11*. Growth kinetics in liquid culture of indicated strains after shift to 18°C.

**Extended Data Figure 8.**
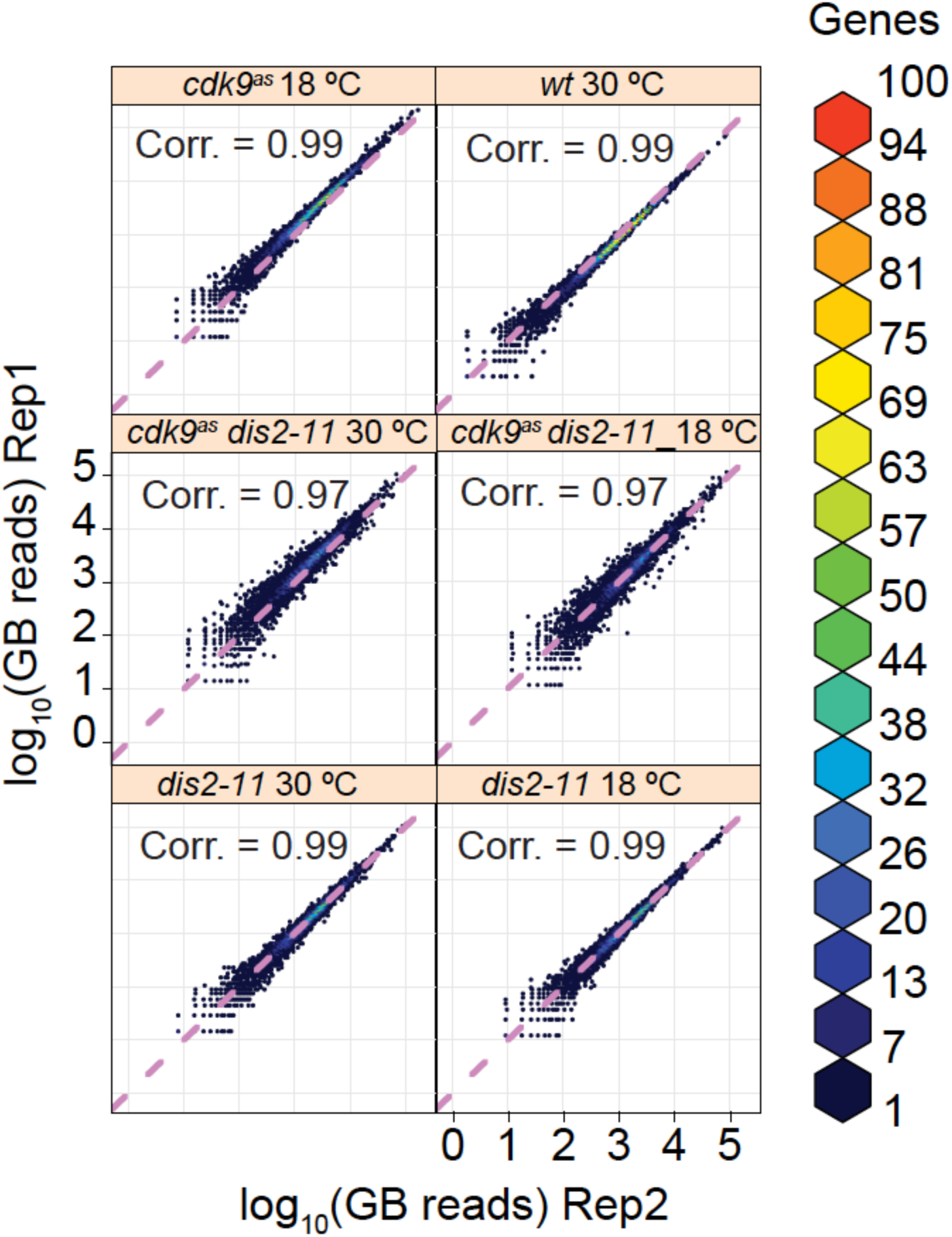
PRO-seq experiments are reproducible. Scatter plots comparing PRO-seq libraries from two biological replicates for each experiment. Values represent log_10_ (normalized reads) within the gene body (TSS + 200 bp to CPS) of all filtered genes (n = 3330). Colors indicate the number of genes represented by each point. Spike-in based normalization should center scatter about the diagonal line x = y (magenta, dotted). Correlation values represent Spearman’s rank correlation.

**Extended Data Figure 9.**
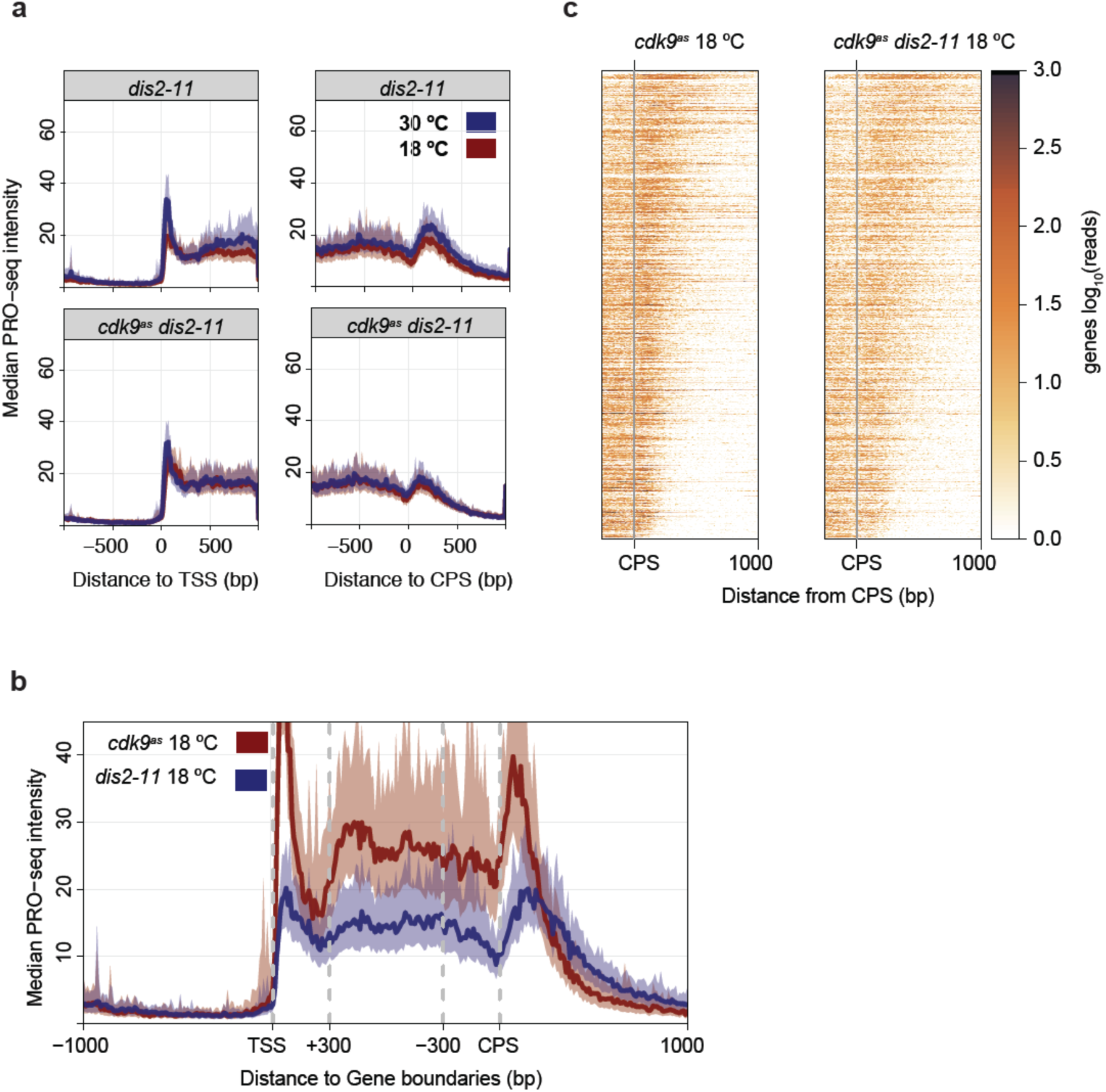
The *dis2-11* mutation affects global transcription properties independent of temperature. **a**, Comparison of composite PRO-seq profiles of the *dis2-11* mutant alone (top panels) or *cdk9*^*as*^ *dis2-11* (bottom panels) at 18°C and 30°C. Profiles are either centered on the TSS (left) or CPS (right). Shaded areas on composite profiles represent the 12.5 and 87.5% quantiles at each position. Each panel represents data from filtered genes that are at least 1 kb from neighboring genes on the same strand (n = 919). **b**, Composite PRO-seq profiles comparing a *dis2*^+^ strain (*cdk9*^*as*^) with the *dis2-11* strain, both at 18°C. Genes were scaled to a common length by fixing the middle gene body (TSS + 300 bp to CPS – 300 bp) region to 60 windows. **c**, Heat maps of spike-in normalized PRO-seq signal (log_10_) within 10 bp windows relative to the CPS (-250 to +1000) for *cdk9*^*as*^ (left) and *cdk9*^*as*^*dis2-11* (right) strains at 18°C. Genes were ranked by decreasing T.E.I. in *cdk9*^*as*^ *dis2-11* at 18°C, a measure of termination-window size.

**Extended Data Table 1.**
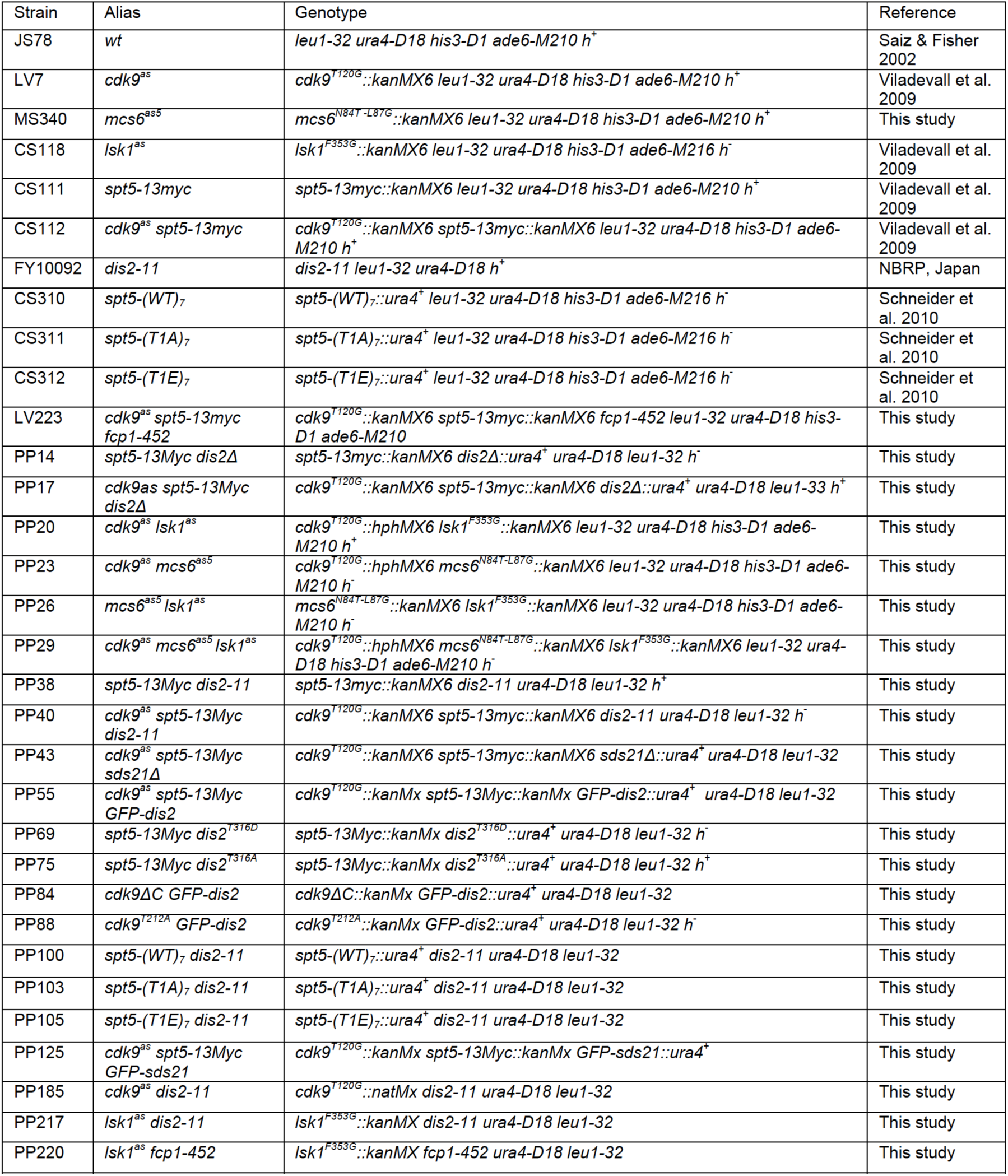
Yeast Strains.

| Strain | Alias | Genotype | Reference |
| --- | --- | --- | --- |
| JS78 | wt | *leu1-32 ura4-D18 his3-D1 ade6-M210 h<sup>+</sup>* | Saiz & Fisher 2002 |
| LV7 | *cdk9<sup>as</sup>* | *cdk9<sup>T120G</sup>::kanMX6 leu1-32 ura4-D18 his3-D1 ade6-M210 h<sup>+</sup>* | Viladevall et al. 2009 |
| MS340 | *mcs6<sup>as5</sup>* | *mcs6<sup>N84T-L87G</sup>::kanMX6 leu1-32 ura4-D18 his3-D1 ade6-M210 h<sup>+</sup>* | This study |
| CS118 | *lsk1<sup>as</sup>* | *lsk1<sup>F353G</sup>::kanMX6 leu1-32 ura4-D18 his3-D1 ade6-M216 h<sup>-</sup>* | Viladevall et al. 2009 |
| CS111 | *spt5-13myc* | *spt5-13myc::kanMX6 leu1-32 ura4-D18 his3-D1 ade6-M210 h<sup>+</sup>* | Viladevall et al. 2009 |
| CS112 | *cdk9<sup>as</sup> spt5-13myc* | *cdk9<sup>T120G</sup>::kanMX6 spt5-13myc::kanMX6 leu1-32 ura4-D18 his3-D1 ade6-M210 h<sup>+</sup>* | Viladevall et al. 2009 |
| FY10092 | *dis2-11* | *dis2-11 leu1-32 ura4-D18 h<sup>+</sup>* | NBRP, Japan |
| CS310 | *spt5-(WT)<sub>7</sub>* | *spt5-(WT)<sub>7</sub>::ura4<sup>+</sup> leu1-32 ura4-D18 his3-D1 ade6-M216 h<sup>-</sup>* | Schneider et al. 2010 |
| CS311 | *spt5-(T1A)<sub>7</sub>* | *spt5-(T1A)<sub>7</sub>::ura4<sup>+</sup> leu1-32 ura4-D18 his3-D1 ade6-M216 h<sup>-</sup>* | Schneider et al. 2010 |
| CS312 | *spt5-(T1E)<sub>7</sub>* | *spt5-(T1E)<sub>7</sub>::ura4<sup>+</sup> leu1-32 ura4-D18 his3-D1 ade6-M216 h<sup>-</sup>* | Schneider et al. 2010 |
| LV223 | *cdk9<sup>as</sup> spt5-13myc fcp1-452* | *cdk9<sup>T120G</sup>::kanMX6 spt5-13myc::kanMX6 fcp1-452 leu1-32 ura4-D18 his3-D1 ade6-M210* | This study |
| PP14 | *spt5-13Myc dis2Δ* | *spt5-13myc::kanMX6 dis2Δ::ura4<sup>+</sup> ura4-D18 leu1-32 h<sup>-</sup>* | This study |
| PP17 | *cdk9<sup>as</sup> spt5-13Myc dis2Δ* | *cdk9<sup>T120G</sup>::kanMX6 spt5-13myc::kanMX6 dis2Δ::ura4<sup>+</sup> ura4-D18 leu1-32 h<sup>+</sup>* | This study |
| PP20 | *cdk9<sup>as</sup> lsk1<sup>as</sup>* | *cdk9<sup>T120G</sup>::hphMX6 lsk1<sup>F353G</sup>::kanMX6 leu1-32 ura4-D18 his3-D1 ade6-M210 h<sup>+</sup>* | This study |
| PP23 | *cdk9<sup>as</sup> mcs6<sup>as5</sup>* | *cdk9<sup>T120G</sup>::hphMX6 mcs6<sup>N84T-L87G</sup>::kanMX6 leu1-32 ura4-D18 his3-D1 ade6-M210 h<sup>-</sup>* | This study |
| PP26 | *mcs6<sup>as5</sup> lsk1<sup>as</sup>* | *mcs6<sup>N84T-L87G</sup>::kanMX6 lsk1<sup>F353G</sup>::kanMX6 leu1-32 ura4-D18 his3-D1 ade6-M210 h<sup>-</sup>* | This study |
| PP29 | *cdk9<sup>as</sup> mcs6<sup>as5</sup> lsk1<sup>as</sup>* | *cdk9<sup>T120G</sup>::hphMX6 mcs6<sup>N84T-L87G</sup>::kanMX6 lsk1<sup>F353G</sup>::kanMX6 leu1-32 ura4-D18 his3-D1 ade6-M210 h<sup>-</sup>* | This study |
| PP38 | *spt5-13Myc dis2-11* | *spt5-13myc::kanMX6 dis2-11 ura4-D18 leu1-32 h<sup>+</sup>* | This study |
| PP40 | *cdk9<sup>as</sup> spt5-13Myc dis2-11* | *cdk9<sup>T120G</sup>::kanMX6 spt5-13myc::kanMX6 dis2-11 ura4-D18 leu1-32 h<sup>-</sup>* | This study |
| PP43 | *cdk9<sup>as</sup> spt5-13Myc sds21Δ* | *cdk9<sup>T120G</sup>::kanMX6 spt5-13myc::kanMX6 sds21Δ::ura4<sup>+</sup> ura4-D18 leu1-32* | This study |
| PP55 | *cdk9<sup>as</sup> spt5-13Myc GFP-dis2* | *cdk9<sup>T120G</sup>::kanMx spt5-13Myc::kanMx GFP-dis2::ura4<sup>+</sup> ura4-D18 leu1-32* | This study |
| PP69 | *spt5-13Myc dis2<sup>T316D</sup>* | *spt5-13Myc::kanMx dis2<sup>T316D</sup>::ura4<sup>+</sup> ura4-D18 leu1-32 h<sup>-</sup>* | This study |
| PP75 | *spt5-13Myc dis2<sup>T316A</sup>* | *spt5-13Myc::kanMx dis2<sup>T316A</sup>::ura4<sup>+</sup> ura4-D18 leu1-32 h<sup>+</sup>* | This study |
| PP84 | *cdk9ΔC GFP-dis2* | *cdk9ΔC::kanMx GFP-dis2::ura4<sup>+</sup> ura4-D18 leu1-32* | This study |
| PP88 | *cdk9<sup>T212A</sup> GFP-dis2* | *cdk9<sup>T212A</sup>::kanMx GFP-dis2::ura4<sup>+</sup> ura4-D18 leu1-32 h<sup>-</sup>* | This study |
| PP100 | *spt5-(WT)<sub>7</sub> dis2-11* | *spt5-(WT)<sub>7</sub>::ura4<sup>+</sup> dis2-11 ura4-D18 leu1-32* | This study |
| PP103 | *spt5-(T1A)<sub>7</sub> dis2-11* | *spt5-(T1A)<sub>7</sub>::ura4<sup>+</sup> dis2-11 ura4-D18 leu1-32* | This study |
| PP105 | *spt5-(T1E)<sub>7</sub> dis2-11* | *spt5-(T1E)<sub>7</sub>::ura4<sup>+</sup> dis2-11 ura4-D18 leu1-32* | This study |
| PP125 | *cdk9<sup>as</sup> spt5-13Myc GFP-sds21* | *cdk9<sup>T120G</sup>::kanMx spt5-13Myc::kanMx GFP-sds21::ura4<sup>+</sup>* | This study |
| PP185 | *cdk9<sup>as</sup> dis2-11* | *cdk9<sup>T120G</sup>::natMx dis2-11 ura4-D18 leu1-32* | This study |
| PP217 | *lsk1<sup>as</sup> dis2-11* | *lsk1<sup>F353G</sup>::kanMX dis2-11 ura4-D18 leu1-32* | This study |
| PP220 | *lsk1<sup>as</sup> fcp1-452* | *lsk1<sup>F353G</sup>::kanMX fcp1-452 ura4-D18 leu1-32* | This study |

